# A single-cell massively parallel reporter assay detects cell type specific cis-regulatory activity

**DOI:** 10.1101/2021.11.11.468308

**Authors:** Siqi Zhao, Clarice KY Hong, Connie A Myers, David M Granas, Michael A White, Joseph C Corbo, Barak A Cohen

## Abstract

Massively parallel reporter gene assays are key tools in regulatory genomics, but cannot be used to identify cell-type specific regulatory elements without performing assays serially across different cell types. To address this problem, we developed a single-cell massively parallel reporter assay (scMPRA) to measure the activity of libraries of *cis-regulatory* sequences (CRSs) across multiple cell-types simultaneously. We assayed a library of core promoters in a mixture of HEK293 and K562 cells and showed that scMPRA is a reproducible, highly parallel, single-cell reporter gene assay that detects cell-type specific *cis*-regulatory activity. We then measured a library of promoter variants across multiple cell types in *ex vivo* mouse retinas and showed that subtle genetic variants can produce cell-type specific effects on *cis*-regulatory activity. We anticipate that scMPRA will be widely applicable for studying the role of CRSs across diverse cell types.

## Introduction

The majority of heritable variation for human diseases maps to the non-coding portions of the genome^1–6^. This observation has led to the hypothesis that genetic variation in the *cis*-regulatory sequences (CRSs) that control gene expression underlies a large fraction of disease burden^7–10^. Because many CRSs function only in specific cell types^11^, there is intense interest in high-throughput assays that can measure the effects of cell-type-specific CRSs and their genetic variants.

Massively Parallel Reporter Assays (MPRAs) are one family of techniques that allow investigators to assay libraries of CRSs and their non-coding variants *en masse^12–18^*. In an MPRA experiment, every CRS drives a reporter gene carrying a unique DNA barcode in its 3’ UTR, which allows investigators to quantify the activity of each CRS by the ratio of its barcode abundances in the output RNA and input DNA. This approach allows investigators to identify new CRSs, assay the effects of non-coding variants, and discover general rules governing the functions of CRSs^12,19–23^. One limitation of MPRAs is that they are generally performed in monocultures, or as bulk assays across the cell types of a tissue. Performing cell-type specific MPRAs in tissues will require methods to simultaneously readout reporter gene activities and cell type information in heterogeneous pools of cells.

To address this problem, we developed scMPRA, a procedure that combines single-cell RNA sequencing with MPRA. scMPRA simultaneously measures the activities of reporter genes in single cells and the identities of those cells using their single-cell transcriptomes. The key component of scMPRA is a two-level barcoding scheme that allows us to measure the copy number of all reporter genes present in a single cell from mRNA alone. A specific barcode marks each CRS of interest (CRS barcode, “cBC”) and a second random barcode (rBC) acts as a proxy for DNA copy number of reporter genes in single cells (**Fig. 1a**). The critical aspect of the rBC is that it is complex enough to ensure that the probability of the same cBC-rBC appearing in the same cell more than once is vanishingly small. In this regime, the number of different cBC-rBC pairs in a single cell becomes an effective proxy for the copy number of a CRS in that cell. Even if a cell carries reporter genes for multiple different CRSs, and each of those reporter genes is at a different copy number, we can still normalize each reporter gene in each individual cell to its plasmid copy number. With this barcoding scheme, we can measure the activity of many CRSs with different input abundances at single-cell resolution, which allows us to measure the activity of CRSs simultaneously across different populations of cells.

**Fig. 1.**
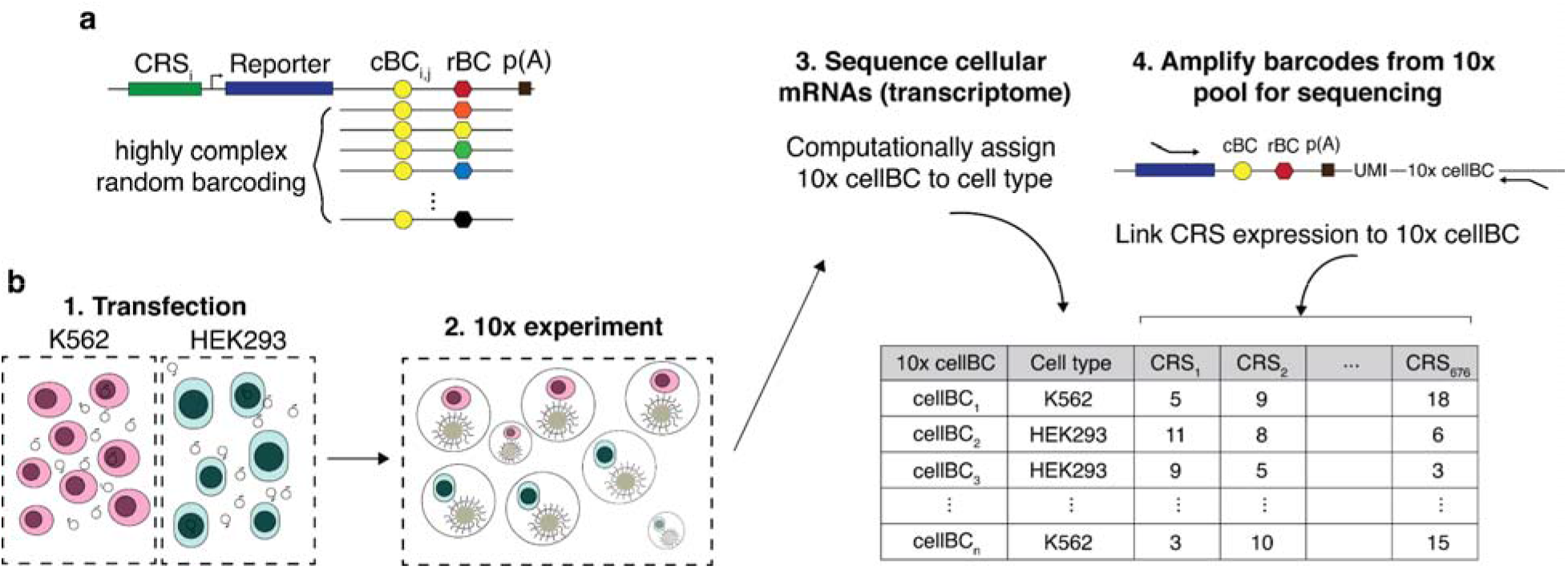
scMPRA measures CRS at single-cell resolution. (**a**) Each CRS reporter construct is barcoded with a cBC that specifi**es** the identity of the CRS, and a highly complex rBC. The complexity of the cBC-rBC pair ensures that the probability of identical plasmids being introduced into the same cell is extremely low. (**b**) Experimental overview for scMPRA using the mixed-cell experiment as an example. K562 cells and HEK293 cells are transfected with the double-barcoded core promoter library. After 24 hours, cells were harvested and mixed for 10x scRNA-seq. Cell identities were obtained by sequencing the transcriptome, and single-cell expression of CRSs were obtained by quantifying the barcodes. The cell identity and CRSs expression (as measured by the cBC-rBC abundances) were linked by the shared 10x cell barcodes.

## Results

### scMPRA enables single-cell measurement of CRS activity

As a proof of principle, we first used scMPRA to test whether different classes of core promoters show different activities in different cell types. Core promoters are the non-coding sequences that surround transcription start sites, where general cofactors interact with RNA polymerase II^24,25^. Core promoters are divided into different classes by the functions of their host genes (housekeeping vs developmental), as well as by the sequence motifs they contain (TATA-box, downstream promoter element (DPE), and CpG islands). We selected 676 core promoters that we previously tested^24,25^ and cloned them into a double-barcoded MPRA library. Given the complexity of the library (>1×10^7^ unique cBC-rBC pairs), we calculated that the probability of plasmids with the same cBC-rBC pair occurring in the same cell is less than 0.01 with our transfection protocols. Given this low likelihood, the number of rBC per cBC in a cell represents the copy number of a CRS in that cell. Knowing the copy number of CRSs in single cells allows us to normalize reporter gene expression from each CRS to its copy number in individual cells.

We performed a cell mixing experiment to test whether scMPRA could measure cell type specific expression of reporter genes. We transfected K562 (chronic myelogenous leukemia) and HEK293 (human embryonic kidney) cells, and performed scMPRA on a 1:1 mixture of those cell lines (**Fig. 1b**). The mRNA from single cells was captured, converted to cDNA, and sequenced. The resulting cBC-rBC abundances and transcriptome of each single cell are linked by their shared 10x cell barcode.

We recovered a total of 3112 cells (97%) that could be unambiguously assigned to one of the two cell types (**Fig. 2a, Supplementary Fig. 1a,b**) and computed the mean expression of each core promoter in the library in each cell type (**Methods**). The measurements were reproducible in both cell types (K562: Pearson’s R = 0.89, Spearman’s ρ = 0.57, HEK293: Pearson’ R = 0.96, Spearman’s ρ = 0.92) (**Figs. 2b,c, Supplementary Table 1**), and we obtained measurements for 99.5% of core promoters in K562 cells and 100% in HEK293 cells, highlighting the efficiency of scMPRA. The median number of cells in which each core promoter was measured was 76 for K562 cells and 287 for HEK293 cells (**Figs. 2d,e**). We also tabulated the number of cBC-rBC pairs in each individual cell and found that the median per cell was 164 in K562 cells and 341 in HEK293 cells (**Supplementary Figs. 1c,d**). On average we detected 10 rBCs per promoter in individual HEK293 cells and 2 rBCs per promoter in K562 cells (**Supplementary Fig. 1e,f**). To validate the scMPRA measurements, we conducted bulk MPRA of the core promoter library in the two cell types separately. Bulk MPRA measurements are not corrected for PCR amplification biases with UMIs, and we found that the bulk measurements correlate well with aggregated single-cell measurements without UMI correction (**Figs. 2f,g)**. That correlation drops with the UMI-corrected single-cell measurements **(Supplementary Figs. 1g,h**), which suggests that bulk measurements may suffer from over counting because of uneven amplification during PCR.

**Fig. 2.**
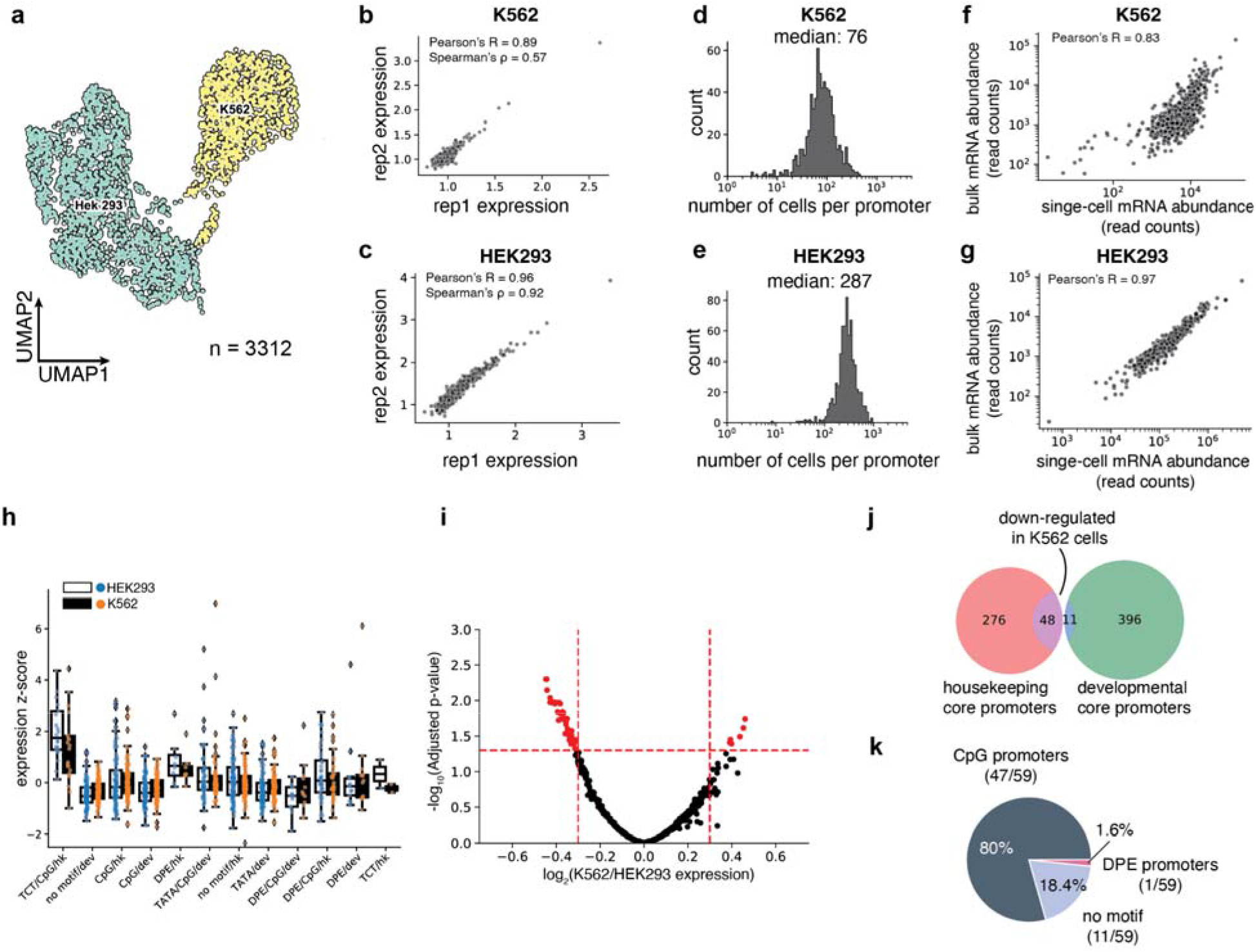
scMPRA detects cell type specific CRS activity. (**a**) UMAP of the transcriptome from the mixed-cell scMPRA experiment. 3312 out of 3417 cells are assigned to either K562 or HEK293 cells. (**b,c**) Reproducibility of replicate measurements of the mean expression from each core promoter in both K562 and HEK293 cells (**d,e**) Histogram of the number of cells in which each core promoter was measured for HEK293 and K562 cells. (**f,g**) Correlations between scMPRA and bulk MPRA using mRNA abundances (cBC counts per cell) to make the two methods comparable. **(h**) Boxplot of the activities of core promoters from different categories in K562 (orange) and HEK293 (blue) cells. The promoter categories are taken from Haberle et al. Because the average expression of all promoters were different between K562 and HEK293, we plotted each category according to its deviation from the average expression (z-score) of all promoters in each cell type (**i**) Volcano plot for differential expression (DE) of core promoters in K562 and HEK293 cells. Significant DE reporters (red dots) have adjusted p-value <0.01 and log-2 fold change greater than 0.3. (**j**) Venn diagram of the functional characterization (housekeeping vs developmental) of down-regulated core promoters in K562 cells. Housekeeping promoters are enriched (p-value = 1.08×10^−11^ from hypergeometric test). (**k**) Pie chart of the sequence features (CpG, DPE, TATA) of down-regulated core promoters. CpG promoters are enriched (p=2.18×10^−6^, from hypergeometric test).

### scMPRA detects cell type specific CRS activity

We asked whether the data allowed us to detect core promoters with differential activity between K562 and HEK293 cells. While different classes of core promoters generally had similar activities in both cell lines (**Fig. 2h**), our differential analysis using DEseq2^26^ identified a small number of promoters (11 out of 669) that are upregulated in K562 cells, and 59 promote**rs** that are downregulated in K562 cells (adjusted p-value < 0.01, log2 fold change > 0.3, **Fig. 2i, Supplementary Table 2**). Among the down-regulated promoters, 48 out of 59 core promote**rs** belong to housekeeping genes (p=1.08×10^-11^**, Fig. 2j**), and 46 out of 59 core promoters are CpG-island-containing core promoters (p=2.18×10^-6^**, Fig. 2k**). This result is not due to differences in the quality of measurements between housekeeping and developmental promoters (**Supplementary Figs. 1i,j**). These results demonstrate the ability of scMPRA to detect CRSs with cell-type specific activities.

### scMPRA detects cell sub-state specific CRS activity

Single-cell studies have revealed heterogeneity in cell states even within isogenic cell types^27–30^. Therefore, we asked if scMPRA can identify CRSs with cell-state specific activity. We repeated scMPRA on K562 cells alone and obtained a total of 5141 cells from two biological replicates. Measurements of each library member were again highly correlated between replicates and agree well with independent bulk measurement (**Supplementary Figs. 2a,b**).

Because the phases of the cell cycle represent distinct cell-states, we asked whether scMPRA could identify reporter genes with differential activity through the cell cycle. We assigned cell cycle phases to each cell using their single cell transcriptome data (**Fig. 3a**) and calculated the mean expression of each reporter gene in different cell cycle phases. We found that most core promoters in our library are upregulated in the G1 phase of the cell cycle, and that some housekeeping promoters are highly expressed through all cell cycle phases (**Fig. 3b, Supplementary Table 3**). We also identified core promoters with different expression dynamics through the cell cycle. For example, we found that the core promoter of *UBA52* remains highly expressed in the S phase, whereas the core promoter of *CXCL10* is lowly expressed throughout (**Supplementary Fig. 2c**). This analysis illustrates the ability of scMPRA to identify CRSs whose expression naturally fluctuates with cellular dynamics.

**Fig. 3.**
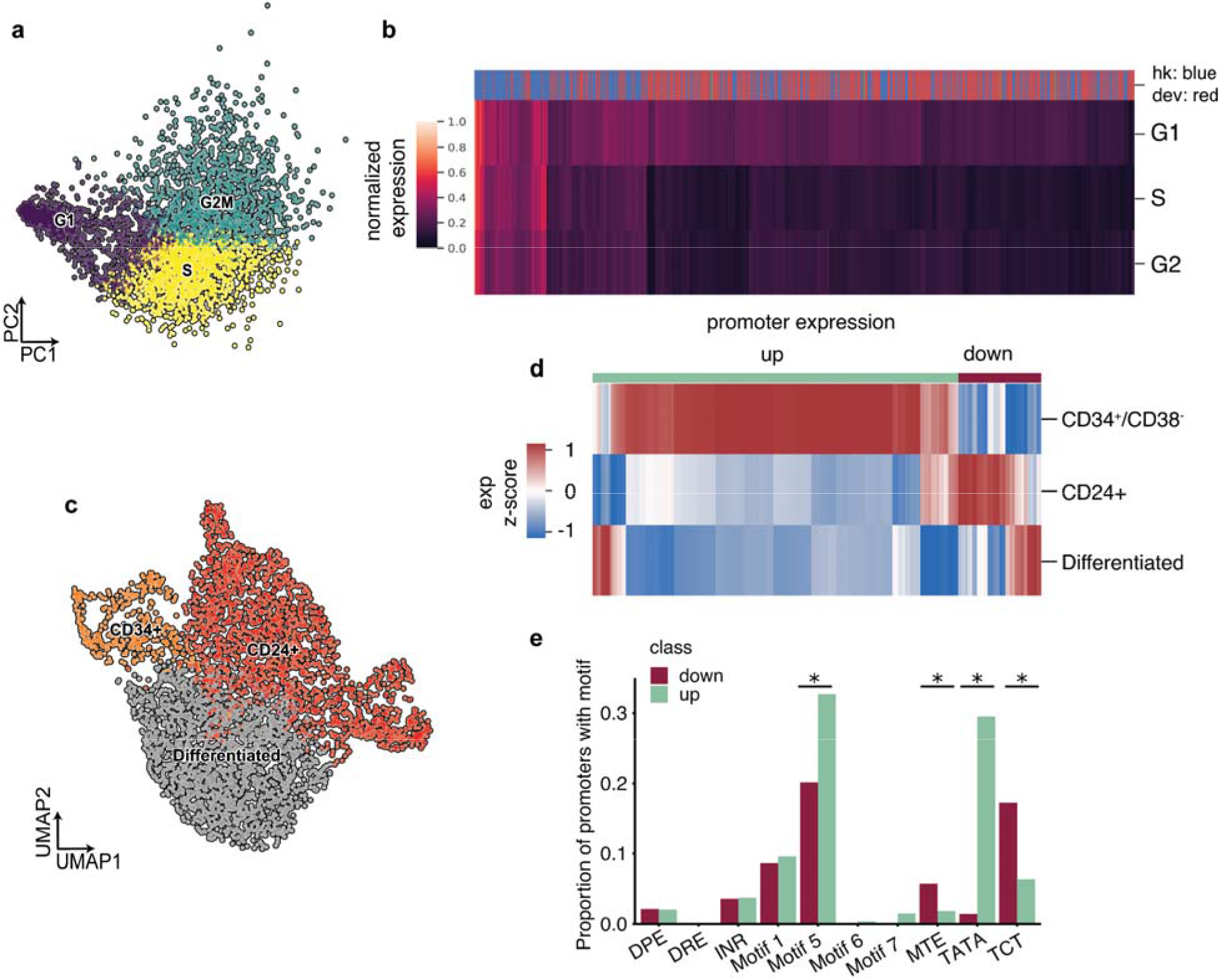
scMPRA detects sub-state-specific CRS activity. (**a**) PCA plot of K562 cells classified by their cell cycle scores. (**b**) Heatmap of core promoter activities in different cell cycle phases (Color bar indicates housekeeping (blue) vs developmental (red) promoters). Core promoter activities have been normalized within each cell cycle phase to highlight the differences between housekeeping and developmental promoters. (**c**) UMAP embedding of K562 cells with high proliferation sub-states (CD34^+^/CD38^-^ and CD24^+^). (**d**) Hierarchical clustering showing two clusters (“up” and “down”) based on expression patterns in the three substates. The promoter activities are plotted as their z-score from the average across cell states to highlight the difference between cell states. (**e**) Proportion of promoters in the up and down clusters that contain the indicated core promoter motif. * represents significant enrichment in one cluster over the other (p < 0.05, Fisher’s exact test).

We then asked whether scMPRA could detect reporter genes with activities that were specific to other cell-states in K562 cells, after normalizing for cell cycle effects. We focused on two specific sub-states that have been reported and experimentally validated for high proliferation rates in K562 cells^31,32^. The first is the CD34^±^/CD38^-^ sub-state that has been identified as a leukemia stem cell subpopulation, and the second is the CD24^+^ sub-state that is linked to selective activation of proliferation genes by bromodomain transcription factors^28,29^. To identify these sub-states in our single-cell transcriptome data, we first regressed out the cell cycle effects and confirmed that the single cell transcriptome data no longer clustered by cell cycle phase (**Supplementary Fig. 2d**). We then identified clusters within K562 cells that have the CD34^±^/CD38^-^ expression signature, or the CD24^+^ signature (**Fig. 3c**). Although the CD34^±^/CD38^-^ cells represent only 9.3% of the cells, scMPRA revealed two distinct classes of core promoters that are upregulated and downregulated in these cells relative to the CD24+ and “differentiated” clusters (**Fig. 3d**). Conversely, the expression patterns of promoters are similar between the CD24^+^ and “differentiated” clusters (**Fig. 3d, Supplementary Table 4**). Motif analysis of the up/down regulated classes of promoters in CD34^+^/CD38^-^ cells showed that different core promoter motifs are enriched in each class, with the TATA box and Motif 5 being enriched in the upregulated class and MTE and TCT motifs being enriched in downregulated class (**Fig. 3e**). This result suggests that differences in core promoter usage might be driving the differences between CD34^±^/CD38^-^ and the other clusters. Because the TATA box is mostly found in developmental core promoters, the CD34^±^/CD38^-^ subpopulation likely reflects the more “stem-like” cellular environment in these cells. Our analysis highlights the ability of scMPRA to identify CRSs with differential activity in rare cell populations.

### scMPRA is reproducible and accurate in murine retinas

To demonstrate that scMPRA is applicable in a complex tissue with multiple cell types, we performed experiments in explanted murine retinas. Intact retina from newborn mice can be cultured and transfected *ex vivo*. This system has been useful for bulk MPRA experiments^13,19,33^, but the results from those experiments report the aggregate expression of library members across the cell types of the retina. Performing scMPRA in *ex vivo* retina provided a chance to assay an MPRA library in a living tissue with multiple cell types in their proper three-dimensional organization.

For this analysis we designed a library consisting of two independently synthesized wild-type copies, and 113 variants, of the full-length *Gnb3* promoter (115 library members, **Supplementary Table 5**). We chose the *Gnb3* promoter because it has high activity in photoreceptors and bipolar cells, but lower expression in other interneurons (i.e. amacrine cells) and Müller glia cells. The library contains mutations in the known transcription factor binding sites (TFBSs) in the *Gnb3* promoter as well as mutations that scan across two phylogenetically conserved regions of the promoter (details in Fig. 6 below). We constructed this library of *Gnb3* promoter variants using the double barcoding strategy described above, with one key modification that we now describe.

In the *Gnb3* promoter library we addressed the inability of scMPRA to measure silent library members. In the first iteration of scMPRA, when a library member produces no mRNA barcodes its corresponding plasmid cannot be detected, and thus, a cell containing a silent plasmid is indistinguishable from a cell without a plasmid. This was not a limitation in our experiments with K562 and HEK293 cells because we detected expressed mRNAs from every member of the library. However, to avoid this potential problem in our retina experiments, we included an additional cassette on the *Gnb3* promoter library that allows us to detect the presence of plasmids carrying silent promoter variants. We included a cassette in which the U6 promoter drives the expression of a second copy of the cBC coupled to the 10x Capture Sequence (**Fig. 4a**). The U6 promoter drives strong RNA Polymerase III-dependent transcription, and is independent of the activity of the *Gnb3* promoter. The Capture Sequence is a specific sequence that is typically used to identify gRNAs in Perturb-seq experiments^34^, but we use it here to isolate U6 expressed cBCs (**Fig. 4b**). When a cell contains a U6 cBC without the corresponding *Gnb3* promoter cBC, it indicates the presence of a silent library member.

**Fig. 4.**
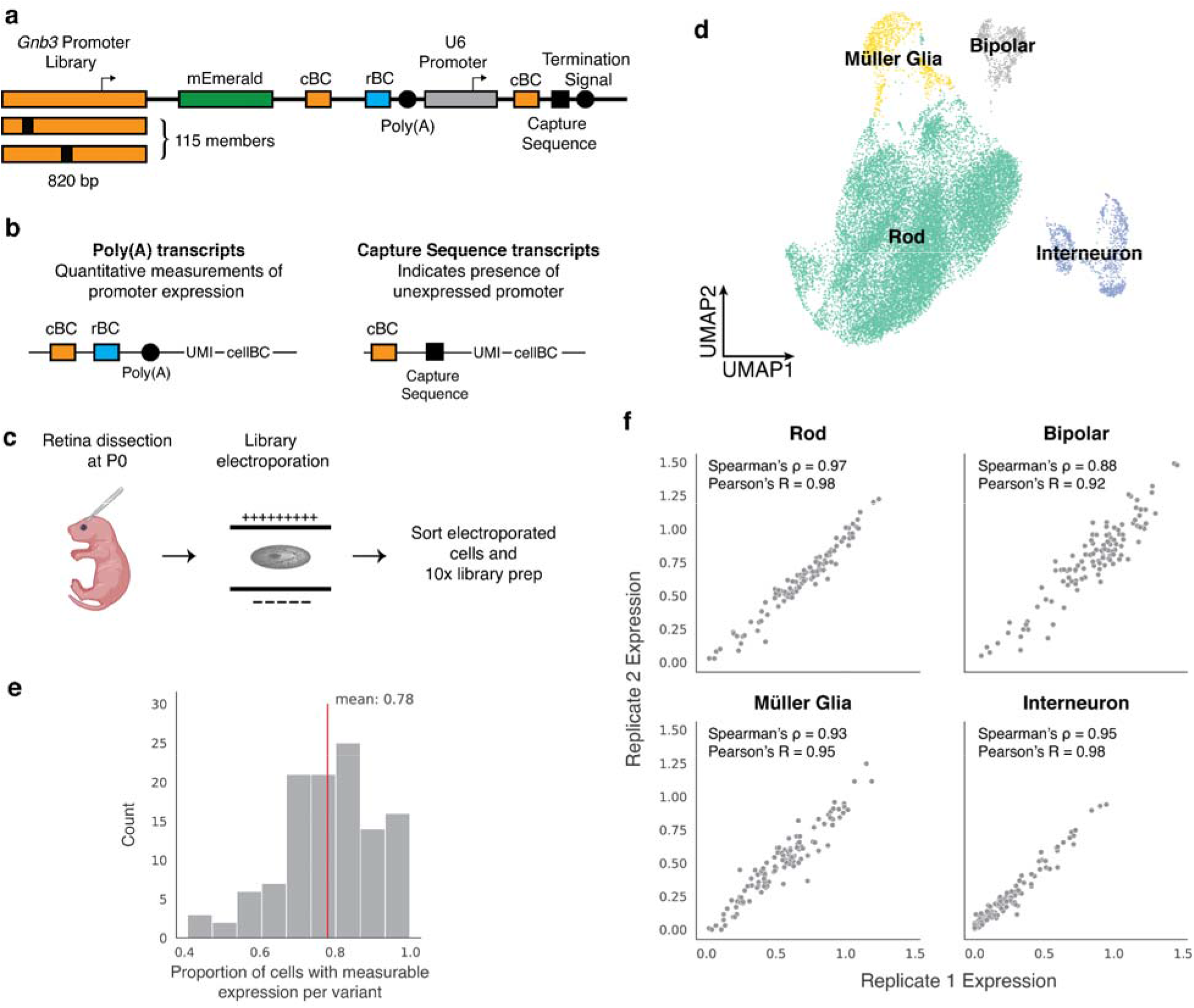
scMPRA design and workflow in mouse retina. (**a**) Schematic of *Gnb3* promoter library constructs. In addition to the cBC and rBC barcodes, the *Gnb3* promoter library contains an additional cassette in which the constitutive U6 promoter expresses a second copy of the cBC with a capture sequence for isolating these transcripts on gel beads. (**b**) Two different types of transcripts produced from the *Gnb3* promoter library to measure promoter expression and detect unexpressed promoters respectively. The two types of transcripts originating from the same cell share the same 10x cell barcodes (**c**) Experimental workflow for scMPRA in *ex vivo* mouse retinas. (**d**) UMAP of all cells measured in scMPRA with four major cell types identified. (**e**) For each *Gnb3* variant in the library, we determined the proportion of cells that contain barcoded poly(A) transcripts out of all the cells that contained the variant. (**f**) Reproducibility of promoter activities between replicates in each of the four major cell types.

We introduced the *Gnb3* promoter variant library into newborn mouse retinas and assessed the cell types into which the library entered by scRNA-seq (**Fig. 4c**). We obtained a total of 22,161 cells from two replicate experiments with a mean of 22,528 reads per cell and 1,642 genes per cell. The scRNA-seq data showed that we recovered rod photoreceptors (87.3%), bipolar cells (3.5%), interneurons (i.e. amacrine cells) (5.2%), and Müller glia cells (3.9%) (**Fig. 4d, Supplementary Figure 3 a,b**).

We then computed the expression of each *Gnb3* promoter variant in each cell type by sequencing the *Gnb3*-expressed barcodes and the U6 barcodes from single cells. Cells with U6-expressed cBC counts, but no *Gnb3*-expressed cBC counts, represented cells in which that promoter variant was silent. On average, *Gnb3* promoter variants were silent in 22% of cells, but this number varied widely (**Fig. 4e**) and was linearly related to the strength of the promoter, with stronger promoters expressing in a larger fraction of cells (**Supplementary Fig. 3c**). Using both the *Gnb3*-driven and U6-driven counts allowed us to compute the average expression of a promoter variant across all the cells of a given cell type, while still accounting for cells in which that promoter variant is silent (**Methods**).

Replicate measurements of the *Gnb3* promoter variant library were reproducible in all four cell types (**Fig. 4f, Supplementary Table 6**). Reproducibility was highest in rod cells (Spearman’s ρ: 0.97, Pearson’s R: 0.98) because rod cells are the most abundant cell type in the mouse retina. The reproducibility was slightly lower in the rarer cell types (bipolar cells: Spearman’s ρ: 0.88, Pearson’s R: 0.92, Müller glia: Spearman’s ρ: 0.93, Pearson’s R: 0.95, and interneurons: Spearman’s ρ: 0.95, Pearson’s R: 0.98), but remained high enough to assess the expression of individual library members. We determined how reproducibility scales with the number of cells in scMPRA by subsampling the expression data. The minimum number of cells required for reproducible measurements (Spearman’s ρ > 0.75) of mean reporter gene levels is 75 cells (**Supplementary Fig. 3d**). Our results show that scMPRA works well for measuring reporter gene levels across cell types in complex tissues using small numbers of cells.

Two additional observations suggest that scMPRA measurements are accurate in *ex vivo* retinas. First, the expression of the wild type *Gnb3* reporter, as well as the average expression of all *Gnb3* promoter variants, correlates with endogenous *Gnb3* expression in the corresponding scRNA-seq data (**Fig. 5a,b**). Second, our scMPRA data reproduced the known effect of a cell type specific *Gnb3* promoter variant. Murphy *et al*.^42^ showed that altering two of the K50 homeobox sites in the *Gnb3* promoter to Q50 sites reduces expression in bipolar cells while leaving expression in rod cells relatively unaffected. We observed the same reduction in bipolar cells when compared with rod cells for this same mutant (**Fig. 5c**). In addition, scMPRA also revealed that this mutant shows increased activity in Müller glia and interneurons. Taken together, these observations demonstrate that scMPRA is reproducible and accurate when applied to cell types in a complex tissue.

**Fig. 5.**
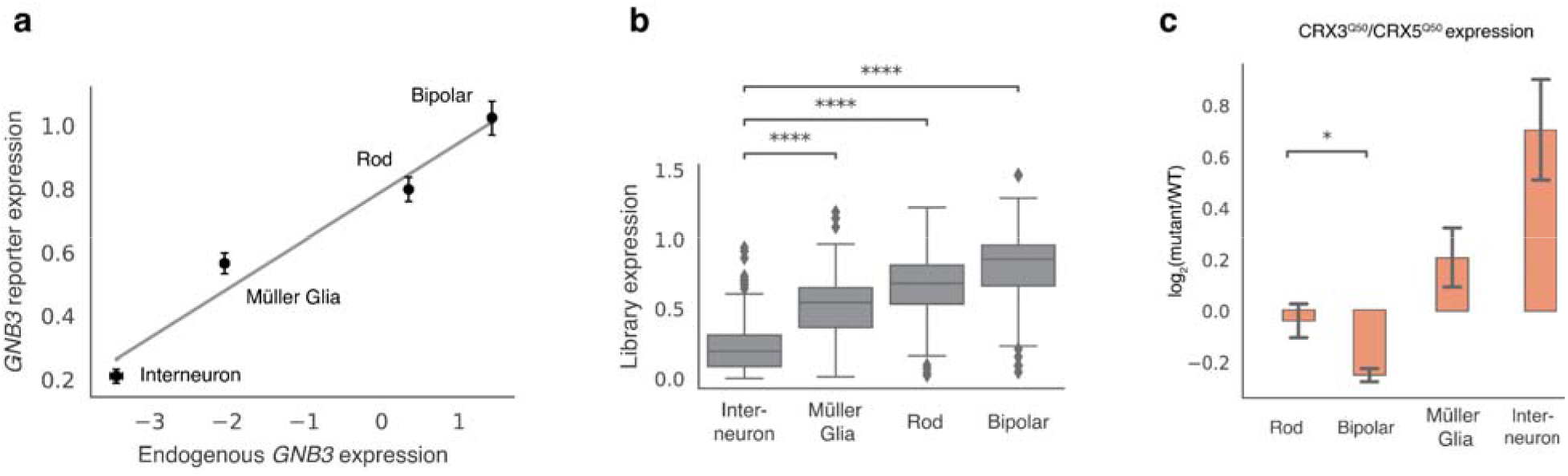
scMPRA recapitulates *Gnb3* expression patterns. (**a**) The expression of the wild-type *Gnb3* promoter in scMPRA reflects endogenous expression levels of *Gnb3* in the respective cell types (Error bar denotes 95% C.I from two biological replicates). (**b**) The expression of the entire *Gnb3* library in different cell types also follows endogenous *Gnb3* expression (****: p-value < 0.0001 for Mann-Whitney U test). (**c**) scMPRA recapitulates the effects of a known *Gnb3* variant, where the CRX3^Q50^/CRX5^Q50^ variant reduces expression in bipolar cells specifically (*: p-value < 0.05 for Welch’s t-test).

### scMPRA reveals cell-type specific promoter variants

The *Gnb3* library was designed to probe components of the promoter including five binding sites for K50-type homeodomain TFs, an E-box binding site, and two evolutionarily conserved regions (**Fig. 6a**). In this experiment we define the effect of a mutation as its relative fold-change to the WT *Gnb3* promoter in each cell type because the *Gnb3* promoter is expressed at different levels across cell types (**Fig. 5a**). We labeled the homeobox sites as Cone Rod Homeobox (CRX) sites because *CRX* is a K50-homeodomain protein which plays an important role in rods and bipolar cells and is required for *Gnb3* expression. Inactivating mutations in any individual CRX site decreased *Gnb3* reporter expression in bipolar and rod cells, but deletion of either CRX1 or CRX5 also resulted in increased expression in interneurons (**Fig. 6b**). The CRX2 disruption had the largest effect on expression, and mutating the CRX2 site in combination with any other CRX site also caused large reductions in expression in rods and bipolar cells. Single and double swaps of K50 CRX binding sites with Q50 binding sites tended to yield cell-type specific effects, primarily because interneurons displayed larger responses to the Q50 swaps compared with rod and bipolar cells (**Fig. 6c**). Increasing the affinity of CRX sites tended to have mild effects on expression in rods and bipolar cells, but increased expression significantly in interneurons (**Fig. 6d**). The results from modifying CRX sites demonstrated that perturbations to single binding sites can produce cell-type specific effects.

**Fig. 6.**
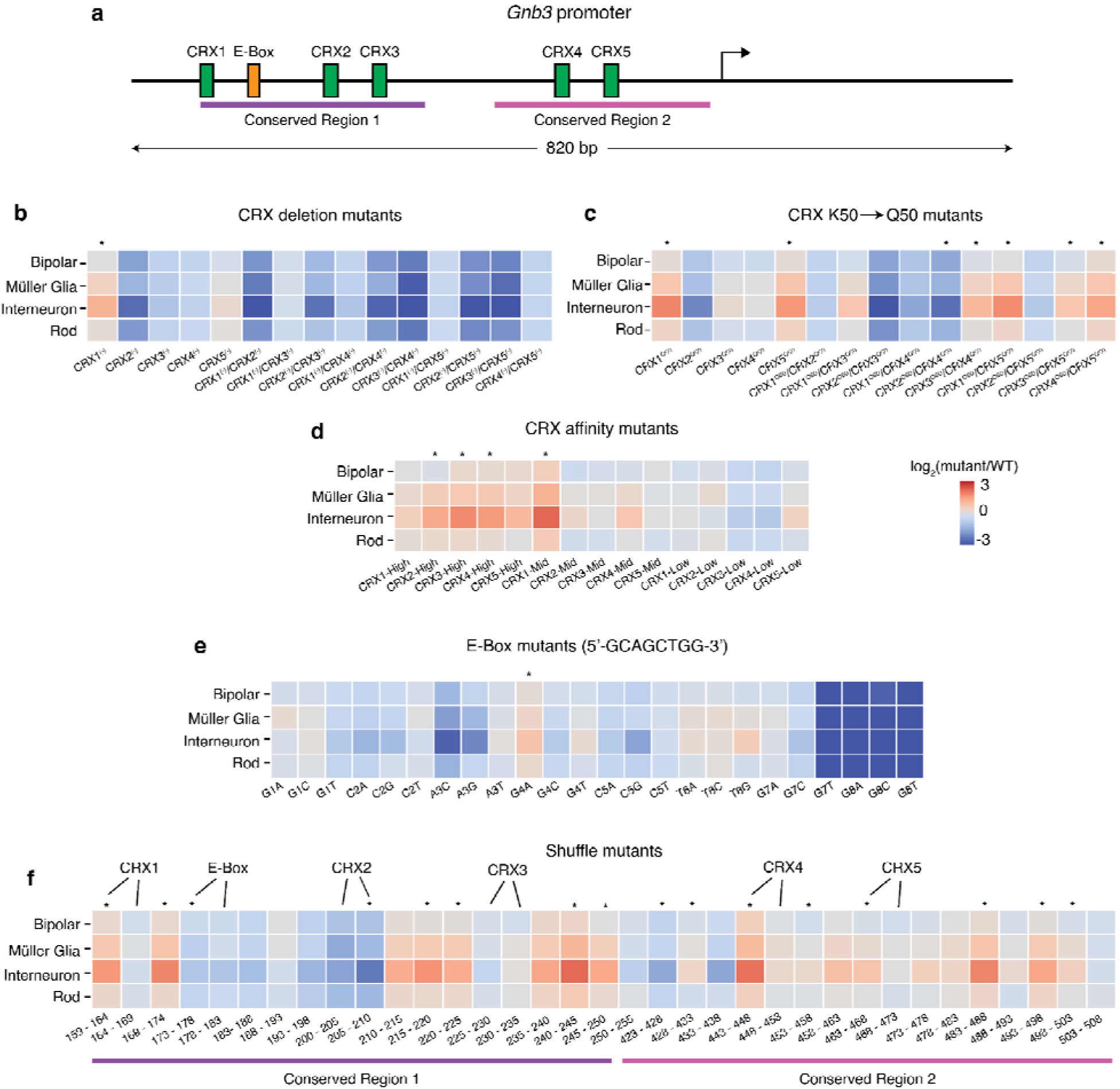
Mutations in the *Gnb3* promoter display cell-type specific effects. (**a**) Schematic of the *Gnb3* promoter showing the location of the five CRX binding sites and the E-box. (**b**) Effects of individual and pairwise deletions of CRX binding sites. (**c**) Effects of individual and pairwise mutations of CRX K50 binding sites to Q50 binding sites. (**d**) Effects of changing CRX binding site affinities. (**e**) Effects of saturation mutagenesis of the E-box. (**f**) Effects of shuffle mutants in conserved regions of the *Gnb3* promoter. Each region was split into 5bp windows and the nucleotides in each window were shuffled. Labels above the heatmap indicate locations where the mutations impact CRX or E-box binding sites. All plots show log2 fold changes of the mutant relative to WT *Gnb3* expression in that cell type. Stars above the plot indicate a significant cell-type specific effect by one-way ANOVA.

We next examined the effects of single nucleotide changes in the E-box binding site (**Fig. 6e**). Helix-Loop-Helix (bHLH) transcription factors, which bind E-box motifs, are critical for the development of multiple retinal cell types^35^. Several single-nucleotide substitutions in the E-box resulted in strong effects on expression, although only one substitution produced significant cell-type specific effects. While the E-box is critical for strong expression of the *Gnb3* promoter, subtle changes to its sequence do not generally result in cell-type specific changes to its activity.

To examine the effects of more severe sequence changes, and to assess the effects of perturbations outside the known TFBSs, we tiled mutations through the two evolutionarily conserved regions shuffling 5 bp at a time (**Fig. 6f**). Mutations in all six TFBSs resulted in cell-type specific changes in expression, but several mutations in the *Gnb3* promoter outside of the known TFBS also resulted in cell-type specific changes in expression. Thus, other information in the *Gnb3* promoter provides important cell-type context for the functioning of the CRX and E-box motifs.

Our analysis of the *Gnb3* promoter shows that single-binding site and single-nucleotide variants can result in cell-type specific changes to *cis*-regulation, and that scMPRA is a powerful tool for identifying these changes across cell-types in mammalian tissues. The *cis*-regulatory logic of the *Gnb3* promoter keeps it expressed at high levels in rods and bipolar cells in the early postnatal period, and at much lower levels in interneurons, which we speculate is why most cell-type specific perturbations result in effects of different sizes in interneurons when compared with rods and bipolar cells.

## Conclusions

We have presented a single-cell MPRA method to measure the cell-type and cell-state specific effects of CRSs. We demonstrated that scMPRA detects cell-type specific reporter gene activity in a mixed population of cells as well as in living retinas, and cell-state specific activity in isogenic K562 cells. The assay is reproducible and reports accurate mean levels of reporter gene activity in as few as 75 cells in a complex tissue. New methods that increase the number of single cells measured per experiment^36^ will increase the size of libraries that can be assayed by scMPRA. The dynamic range was relatively small in this study (8-fold between the strongest and weakest *Gnb3* variants), which may reflect the activity of these specific sequences, but may also arise from the low efficiency of mRNA capture in single cells. scMPRA will therefore benefit from continuing improvements of methods to capture and recover mRNA from single cells.

With the burgeoning of Adeno-associated viral delivery systems^37–41^, we anticipate that scMPRA will be widely used to study *cis*-regulatory effects in a variety of complex tissues. Given the hypothesis that non-coding variants with cell-type specific effects underlie a large fraction of human disease, an important application of scMPRA will be to identify genetic variants with cell-type specific *cis*-regulatory effects.

## Methods

### Cell Culture

K562 cells were cultured in Iscove’s Modified Dulbecco’s Medium (IMDM) + 10% Fetal Bovine Serum (FBS) + 1% non-essential amino acids + 1% pen/strep at 37 □ with 5% CO_2_. HEK293 cells were cultured in Eagle’s Minimum Essential Medium (EMEM) + 10% Fetal Bovine Serum (FBS) + 1% pen/strep at 37 with 5% CO_2_.

### Core Promoter Library Cloning

We developed a two-level barcoding strategy to enable single-cell normalization of plasmid copy number. We applied this strategy to a library of core promoters we previously tested by bulk MPRA^24^. That core promoter library contains 676 core promoters, each with a length of 133bp. The library cloning was done in three steps: First, we synthesized a library of 676 core promoters each barcoded with 10 different cBCs and cloned this library into a backbone^24^. In a second step, a dsRed fluorescent reporter cassette was cloned between each core promoter and its associated cBCs as described^24^. Thirdly, we modified this library for scMPRA by adding random barcodes downstream of the cBCs, but upstream of the polyA site.

To add the random barcodes (rBCs) we synthesized a single-stranded 90 bp DNA oligonucleotide (oligo) containing a 25 bp random sequence (the rBC), a restriction site, and 30 bp homology to the library vector on each side of the rBC region. We used NEBuilder^®^ HiFi DNA Assembly Master Mix (E2621) to clone this oligo into the core promoter library. 4μg of the plasmid library was split into four reactions and digested with 2μl of SalI for 1.5 hours at 37□. The digested product was run at 100V for 2 hours on a 0.7% agarose gel. The correct size band was cut and purified with the Monarch Gel Extraction Kit (New England BioLabs T1020L). The insert single-stranded DNA was diluted to 1 μM with H2O. Three assembly reactions were pooled together, each reaction containing 100 ng of digested library backbone, 1μM of insert DNA, 1 μl of NEBuffer 2, 10 μl of 2X Hifi assembly mix, and H2O up to 20 μl. The reaction was incubated at 50°C for 1 hour. The assembled product was purified with the Monarch PCR&DNA Cleanup kit (New England BioLabs T1030L) and eluted in 12 μl of H2O.

The assembled plasmid was transformed using Gene Pulser Xcell Electroporation Systems by electroporation (BIO-RAD 1652661) into 50 μl of ElectroMax DH10B electrocompetent cells (Invitrogen 18290015) with 1 μl of assembled product at 2 kV, 2000 Ω, 25 nF, with 1 mm gap. 950 μl of SOC medium (Invitrogen 15544034) was added to the cuvette and then transferred to a 15 ml Falcon tube. Two transformations were performed, and each tube was incubated at 37□ for 1 hour on a rotator with 300 rpm. The culture was then added to pre-warmed 150 μl LB/Amp medium and grown overnight at 37□. 1 μl of the culture was also diluted 1:100 and 50 μl of the diluted cultured was plated on a LB agar plate to estimate the transformation efficiency. For the core promoter library, we prepared DNA from more than 4×10^8^ colonies. Shallow sequencing of this library (below) showed that the majority of library members encoded unique cBC-rBC combinations.

### *Gnb3* Promoter Variant Library Design and Cloning

The *Gnb3* library was designed to probe components of the promoter including five binding sites for K50-type homeodomain TFs, an E-box binding site, and two evolutionarily conserved regions. We labeled the K50 homeobox sites as Cone Rod Homeobox (CRX) sites because CRX is a K50-homeodomain protein required for *Gnb3* expression and a key-lineage determining factor in retina, even though other K50-type homeobox proteins are also expressed in retinas. To test whether the disruption of CRX sites in the *Gnb3* promoter has cell-type specific effect, we made the following three types of mutations: (1) All individual and pairwise deletions of the CRX binding sites by mutating ^42^ the CRX sites to 5’-CTACTCCC-3’. (2) All individual and pairwise mutations of CRX binding sites from K50 homebox to Q50 homeobox motifs: 5’-CTAATTAC-3’. (3) All individual mutations of CRX binding sites to high (5’-CTAATCCC-3’), medium (5’-CTAAGCCC-3’) and low affinity (5’-CTTATCCC-3’) K50 homeobox sites ^22^. Our unpublished data suggested that the E-box is important for the Gnb3 promoter activity and E-box motif is bound by many neuronal specific TFs ^35^, hence we mutated each base pair in the E-box to every other base pair and made pairwise mutations of the two core base pairs in the E-box motif. Lastly, we took an unbiased approach to screen for potential cell-type-specific mutations by shuffling mutations across the two conserved regions in the *Gnb3* promoter. Each conserved region was tiled into 5bp windows and the nucleotides within each window were shuffled. All library sequences and the corresponding cBCs can be found in Supplementary Table 5.

The library of *Gnb3* promoter variants was constructed in four steps. In the first step, we cloned the *Gnb3* promoter variant library into the core promoter library vector backbone. We ordered double-stranded DNA fragments from Integrated DNA Technologies (Coralville, Iowa) encoding the varying part of the (520 bp) *Gnb3* promoter and 113 promoter variants. The wild type *Gnb3* promoter sequence was included twice, each time fused to a different cBC. The DNA fragments were manually pooled and cloned together as a library. In the second step, we cloned the remaining *Gnb3* promoter (300bp) and an mEmerald reporter cassette between the *Gnb3* promoter variants and the first cBC copy using HiFi assembly. In the third step, we used NEB HiFi DNA Assembly Master Mix (New England Biolabs E2621) to insert the U6 promoter between the two copies of the cBCs where it drives expression of the downstream copy of the cBC. In the fourth step, we introduced high-complexity rBCs between the first cBC and the U6 promoter. We synthesized a DNA oligo containing a 25 bp random sequence (the rBC), a restriction site, and 30 bp homology to the library vector on each side of the rBC barcode region. We then used HiFi Assembly to clone the rBC oligos into the *Gnb3* promoter variant library. In this final library each plasmid contains a *Gnb3* promoter variant driving mEmerald with a unique cBC-rBC combination in its 3’ UTR, which is followed by a polyA signal and the U6 promoter driving a second copy of the cBC, a capture sequence, and a termination signal. A total of eight HiFi Assembly reactions were pooled together to increase the library complexity. This library was transformed and amplified in *E. coli* as described above, and DNA was prepared from 2×10^9^ colonies.

### Estimating Library Complexity

To estimate the complexity of the core promoter library, we sequenced the DNA library using a nested PCR-based Illumina library preparation protocol. Briefly, we first used Q5 polymerase (New England BioLabs M0515) to amplify the region containing the two barcodes with SCARED 17 and SCARED P18 (primer sequences can be found in Supplementary Table 8). The total reaction volume was 50μl using 50ng of plasmid library with 2.5 μl of 10uM primer each. After 25 cycles of amplification the product was purified with the Monarch PCR&DNA Cleanup kit (New England BioLabs T1030), and eluted with 20 μl of ddH2O. For the second round of PCR we used the primers SCARED P19 and SCARED P20 in a 25 μl reaction with 0.25 μl product from the previous step. After 10 rounds of amplification the product was purified using the Monarch PCR&DNA Cleanup kit. For the last PCR we added the P5 and P7 Illumina adapters with SCARED P5, SCARED P7 with 10 cycles of amplification in a 25 μl reaction with 2 μl of purified product. This final product was sequenced on an Illumina MiSeq, and we obtained a total of 1,693,933 reads. After filtering out reads without a cBC or rBC of the correct length, we obtained a total of 1,359,176 reads (80% of the total reads) and 99.5% represented unique cBC-rBC pairs. For the *Gnb3* library, we performed shallow sequencing, and obtained a total of 1,939,479 reads. After filtering out reads without correct cBC or rBC, we obtained a total of 1,838,415 reads (94.7% of the total reads). Among the 1,838,415 correct reads, 99.5% represented unique cBC-rBC pairs.

### Estimating the Probability of Identical cBC-rBC Pairs in the Same Cell

We estimated the probability that more than one copy of a plasmid carrying the same cBC-rBC pair would be transfected into the same cell. We call this probability the collision rate. If the library is transfected into *n* cells, and a specific cBC-rBC pair is present at *m* copies in the library, then the expected number of collisions per experiment is given by:

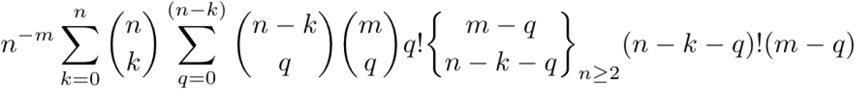

where k denotes the number of cells that received no plasmid, q denotes the number of cells transfected with exactly one plasmid, parentheses denote the binomial coefficient, and bracke**ts** denote the partition function. The above expression was simplified by substituting with the bivariate generating function, and the expected number of collisions per experiment is:

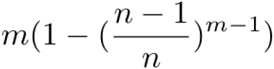

The expected number of collisions per cell (λ) is given by,

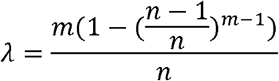

And, assuming collisions are a Poisson process, the probability of at least one collision in a cell is:

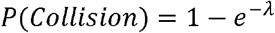

Using this framework, we can estimate the probability of a collision in our experiment. We assume one million cells (*n*) are transfected using 10 ug of plasmid DNA, and that the effective number of plasmids that enter the nucleus is 10% of that input amount (1 ug)^43^1 ug of plasmid DNA is 2.3×10^11^. Thus, the value of *m* in the nucleus is 2.3×10^11^ divided by the number of unique members of the library. This allows us to calculate P(Collision) for a library of any given size. This framework shows that we require a library with 1.6×10^6^ unique members to achieve P(Collision)=0.01. To be 99% sure that a library has at least 1.6×10^6^ unique members requires preparing that library 4.5 times as many independent colonies (7.2×10^6^), assuming a Poisson distributed library. The core promoter library was prepared from 4×10^8^ colonies, 55 times more than required for P(Collision)=0.01, and the *Gnb3* variant library was prepared from 2×10^9^ colonies, 277 times more than required for P(Collision)=0.01.

### Cell Line Transfections

K562 cells were transfected with the core promoter library using electroporation with the Neon transfection system (Invitrogen MPK5000). One million cells were transfected with 2 μg of plasmid DNA (mixed-cell experiment) or 10 μg of plasmid DNA (K562 sub-state experiment), with 3 pulses of 1450 V for 10 ms.

HEK293 cells were transfected with the core promoter library using the Lipofectamine3000 protocol. 4 μl of p3000 reagent, 4μl of Lipofectamine, and OptiMEM were mixed with 2 μg of plasmid DNA to a volume of 250 μl. The lipofectamine reagents and plasmid were mixed and incubated at room temp for 15 minutes and then added dropwise to 1 million cells. We harvested K562 and HEK293 cells 24 hours after transfections for scMPRA.

### *Ex vivo* culturing and transfection of mouse retinas

Retinas from newborn mice were dissected, cultured, and electroporated as described in White et al.^44^ For each replicate experiment, three retinas were electroporated with 0.5 μg/μL of the *Gnb3* promoter variant library, and co-electroporated with 0.5 μg/μL of a plasmid in which the *Rhodopsin* promoter drives the dsRED fluorophore. Electroporated retinas were harvested and dissociated as in Murphy et al.^42^ with modifications as outlined below.

Briefly, three retinas/replicate were washed 3x in cold Hanks’ Balanced Salt Solution (HBSS) (Thermo Fisher) and were then incubated in 400 ul of HBSS containing 0.65 mg papain (Worthington Biochem) for 10 min at 37°C. 600ul of Dulbecco’s Modified Eagle Medium (DMEM) (Thermo Fisher) containing 10% fetal calf serum (FCS) (Gibco) was added and the tissue was gently triturated with P1000 to achieve single cells suspension. 100 units DNase1 (Roche) was added, the cell suspension and incubated an additional 5 min at 37°C. Cells were centrifuged at 400g for 4 min then resuspended in 600 ml of sorting buffer (2.5 mM EDTA, 25 mM HEPES, 1% BSA in HBSS) and passed through a 35um filter and used directly for Fluorescence Activated Cell Sorting (FACS).

Because the majority of cells in murine retinas are rod photoreceptors, we attempted to enrich other cell types using FACS. The co-electroporated Rhodopsin-DsRed construct marks rod cells specifically. Therefore, we used FACS to generate a 1:1 mixture of dsRED^+^ to dsRED^-^ cells from dissociated retinas. This procedure should yield a mix of cells in which rod cells comprise 50% of the total cells. In practice, rod cells still comprised 87% of the cells that were analyzed by scMPRA.

### Bulk MPRA from Cell Lines

For both K562 cells and HEK293 cells, we harvested cells 24 hours after transfection, extracted total mRNA, and performed reverse transcription using the Superscript IV Reverse Transcriptase kit (Invitrogen 18090010). A sequencing library was then constructed using a nested PCR strategy. Briefly, we first used Q5 (New England BioLabs M0515) polymerase to amplify the region containing the cBC-rBC barcodes with SCARED P17 and SCARED P18. The total reaction volume was 50μl with 50ng of backbone and 2.5 μl of 10uM primer each. After 25 cycles of amplification the product was purified with the Monarch PCR&DNA Cleanup kit (New England BioLabs T1030L) and eluted with 20 μl of ddH2O. Sequencing adapters were then added with 2 rounds of PCR, 10 cycles each. 0.25μl of PCR product was used with SCARED P19 and P20, then purified using the Monarch PCR&DNA Cleanup kit (New England BioLabs T1030L). Finally, we used 2 μl of this product with primers SCARED P5 and SCARED P7, and then sequenced the resulting product on an Illumina Mi-seq instrument. The activity of library members was computed as described previously^21^.

### Single-cell RNA-seq for scMPRA

To perform scMPRA we targeted 2000 cells from the HEK/K562 mixed pool per replicate for each mixed cell experiment, 2500 cells per replicate for the K562-only experiment, and 2500 cells (after sorting) per replicate for the retina experiment. The cells were prepared according to the 10x Chromium Single Cell 3’ Feature Barcode Library Kit (PN-1000079) protocol.

Our goal was two-fold: to quantify the cBC-rBC pairs from each single cell and to sequence the cellular mRNAs from those same single cells. We captured all polyadenylated RNAs (barcoded reporter RNAs and cellular mRNAs) from single cells following the manufacturer’s protocol up to the cDNA amplification step.

For the cellular mRNAs (transcriptome), we followed the 10x protocol, using 1/4 of the cDNA library to generate dual-indexed transcriptomes. To quantify the cBC-rBC pairs, we performed separate PCRs using primers specifically targeting the reporter gene to improve barcode recovery efficiencies. Because the 10x protocol only uses 1/4 of the generated cDNA, we separately amplified the barcodes from another 1/4 of the pellet cleanup. We first used Q5 (New England BioLabs M0515) polymerase to amplify the region containing the cBC-rBC pairs with SCARED P17 and SCARED P18 with 10 cycles. The sample was divided equally into eight PCR reactions, each with 50 μl of total volume to reduce possible jackpotting. The product was then purified with the Monarch PCR&DNA Cleanup kit (New England BioLabs T1030L), and eluted with 20 μl of ddH2O. We then added sequencing adapters using an additional two rounds of PCR. The first adapter PCR was performed with SCARED P21 and SCARED PP2 with a total of 10 ng of product from the barcode PCR. Again, we pooled eight PCR reactions, each with 50 μl of total volume and 10 PCR cycles. The PCR product was purified using the Monarch PCR&DNA Cleanup kit (New England BioLabs T1030L). For the last PCR, to add the P5 and P7 Illumina adapters, we used the primers SCARED P45, SCARED PP3 with 10ng of product and pooled eight PCR reactions, each with 50 μl of total volume and 10 PCR cycles.

For the U6 promoter library construction, we followed Step 4 of the 10x feature barcoding library preparation protocol (Chromium Next GEM Single Cell 3’ Reagent Kits v3.1 (Dual Index) CG000316 Rev C).

The transcriptome and barcode libraries were combined and sequenced on the Illumina NextSeq 500 with 28×105 cycles. Read1 was limited to 28bp to avoid sequencing the constant poly(A) sequence.

### scRNA-seq data processing

The single-cell RNAseq data was processed using Cellranger 6.0.1 (https://github.com/10xGenomics/cellranger) and Scanpy 1.8.1^45^ (https://github.com/theislab/scanpy) following the standard pipeline. Briefly, different sequencing runs from the same biological replicate were pooled together and processed with CellRanger 6.1.1; the final output expression matrix was then imported into Scanpy for further processing. We first removed cells with less than 1000 genes, genes that were present in less than three cells, and cells with high counts of mitochondrial genes. Next, we normalized the UMI counts to the total cell UMI counts. The normalized expression matrix was used for clustering and visualization with Scanpy.

### scMPRA Data Processing

For each promoter library, paired-end reads generated from barcoded reporter RNAs were processed with custom scripts that can be found on Github (https://github.com/barakcohenlab/scMPRA). In each paired-end read, Read1 contains a 10x cell barcode and a UMI, while Read2 contains the cBC and rBC sequences. We define a “quad” as a 10x Cell Barcode, UMI, cBC, and rBC originating from the same individual paired-end read. To tabulate the cBC-rBCs we first matched the constant sequences flanking both barcodes, filtering out reads where either barcode was not the correct length. We performed this filtering using a stand-alone program (https://github.com/szhao045/scMPRA_parsingtools). Second, we filtered out incorrect 10x Cell Barcodes based on the CellRanger output barcode list using error-correction with a maximum hamming distance of one. Third, to mitigate the effect of template-switching during the PCR steps, we plotted the rank read depth for each unique quad and identified an “elbow point” at a minimum depth of 1 read for the mixed-cell and the retina experiment, and 10 reads for the K562 alone experiment. We kept all reads above the minimum depth, and also kept a low-depth unique quad if it contained a cBC-rBC matching a high-depth pair with a hamming distance of at most one. Lastly, for the mixed-cell experiment and the K562 cell alone experiment, we removed any cell with less than 100 scMPRA-associated UMIs, since the scMPRA reads from those cells were poorly sampled. For the last step, because the retina experiment contains additional information from the U6 promoter, we did not threshold based on cells. Since U6 promoter data provides information on whether a given cBC in a given cell is sampled well, we removed all unique barcode pairs containing only 1 UMI for a cBC.

### Calculating the Single-Cell Activities of Promoters

Once the high-confidence quads were identified, we computed A, the activity of a promoter in an individual cell using,

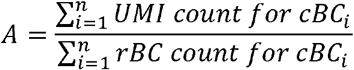

where n is the number of unique cBCs that mark a single promoter in the library, and the UMI and rBC counts are summed over all quads with a given 10x cell barcode. We then compute C, the cell-type specific activity of a promoter as,

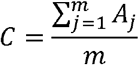

where m is the number of cells in a given cell type, and all 10x cell barcodes assigned to a given cell type are identified from their matched scRNA-seq profiles. For scMPRA data from the retina we modified the equation for cell-type specific activity as follows,

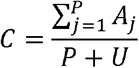

where P is the number of cells of a given cell type in which *Gnb3*-driven cBCs were detected and U is the number of cells of that cell type for which a U6 promoter cBC was detected without detecting any corresponding *Gnb3*-driven cBC. This modification has the effect of adding activities of zero for all cells with U6-driven cBCs that did not express a *Gnb3*-driven cBC.

### Cell cycle analysis

Cell cycle analysis for the scRNA-seq experiment was done with Scanpy 1.8.1 with cell cycle genes^46^. The expression profile of each cell was projected onto a PCA plot based on the list of cell cycle genes using Scanpy.

### Motif analysis

The core promoters were first clustered according to their expression levels in the different cell sub-state populations by hierarchical clustering. We categorized our data into up/down regulated clusters at the first branching point, aiming to preserve the large structure. We then identified core promoter motifs in each promoter according to the parameters in Zabidi et al^47^. using MAST v4.10.0^48^ and plotted the proportion of promoters containing each motif in each promoter class.

### Statistical Analyses

All statistical analyses were done using Python 3.9.6, Numpy 1.12.1^49^, Scipy 1.6.3 and R 4.0.2.

### Ethics

This study was performed in strict accordance with the recommendations in the Guide for the Care and Use of Laboratory Animals of the National Institutes of Health. All of the animals were handled according to protocol # A-3381-01 approved by the Institutional Animal Care and Use Committee of Washington University in St. Louis. Euthanasia of mice was performed according to the recommendations of the American Veterinary Medical Association Guidelines on Euthanasia. Appropriate measures are taken to minimize pain and discomfort to the animals during experimental procedures.

## Data and Code Availability

Next-generation sequencing data that support the findings of the study are available in the Gene Expression Omnibus using accession code GSE188639.

The code that supports the findings of this study is available on Github Repository (https://github.com/barakcohenlab/scMPRA).

## Acknowledgements

We thank the members of the Cohen laboratory for their critical feedback on the manuscript. We thank Jess Hoistington-Lopez and MariaLynn Crosby for assistance with high-throughput sequencing. This work is supported by grants to B.A.C from the National Institutes of Health, R01 GM140711 and R01 GM092910, and by a grant to J.C.C from the National Institutes of Health R01 EY030075.

## Author Contributions

S.Z. and B.A.C. conceived and designed the project. All experiments and analyses were performed by S.Z. with technical contributions from C.K.Y. H. and D.M.G., except for the electroporation and culturing of mouse retinas, which was performed by C.A.M. J.C.C and M.A.W provided critical input into the design of the *Gnb3* promoter variant library. S.Z., C.K.Y.H., and B.A.C. wrote the manuscript with input and feedback from all authors.

## Supplemental Tables

Supplemental Table 1. Mixed Cell Experiment Expression

Supplemental Table 2. Differential expression of the core promoter library between K562 and HEK293 cells

Supplemental Table 3. K562 Cell Cell Cycle Expression

Supplemental Table 4. K562 Cell Cell Sub-state Expression

Supplemental Table 5. Gnb3 Promoter Variant Library

Supplemental Table 6. Gnb3 Library Expression in Retina

Supplemental Table 7. Sequences of Oligonucleotides Used in this Study

## Supplementary Figures

**Supplementary Figure 1.**
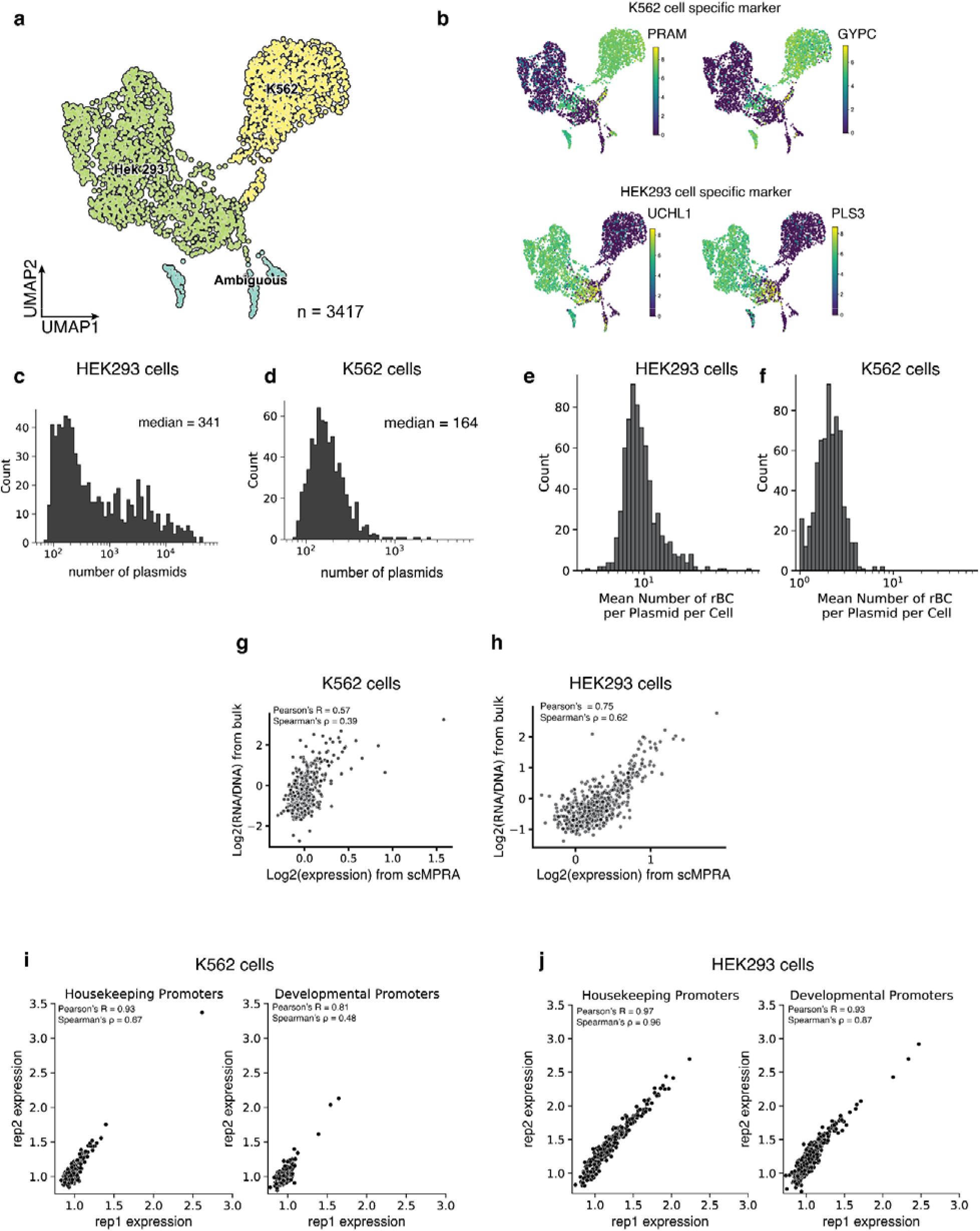
scMPRA measures cell-type specific CRS activity. (**a**) UMAP of the single-cell transcriptome from the mixed-cell experiment. 105 out of 3417 cells (3%) are labeled by both K562 and HEK293 cell genes. (**b**) UMAP of the mixed-cell experiment with cells marked by other representative markers for K562 and HEK293 cell expression. (**c,d**) Histogram of the number of plasmids (unique cBC-rBC pairs) transfected into K562 cells and HEK293 cells. (**e,f**) Histogram of the mean number of rBC per cBC (CRS) per cell for K562 cells and HEK293 cells. (g,h) Correlation of bulk MPRA versus scMPRA where only the scMPRA data has been UMI normalized (**i,j**) Scatterplot of scMPRA reproducibility for housekeeping and developmental promoters in K562 cells and HEK293 cells.

**Supplementary Figure 2.**
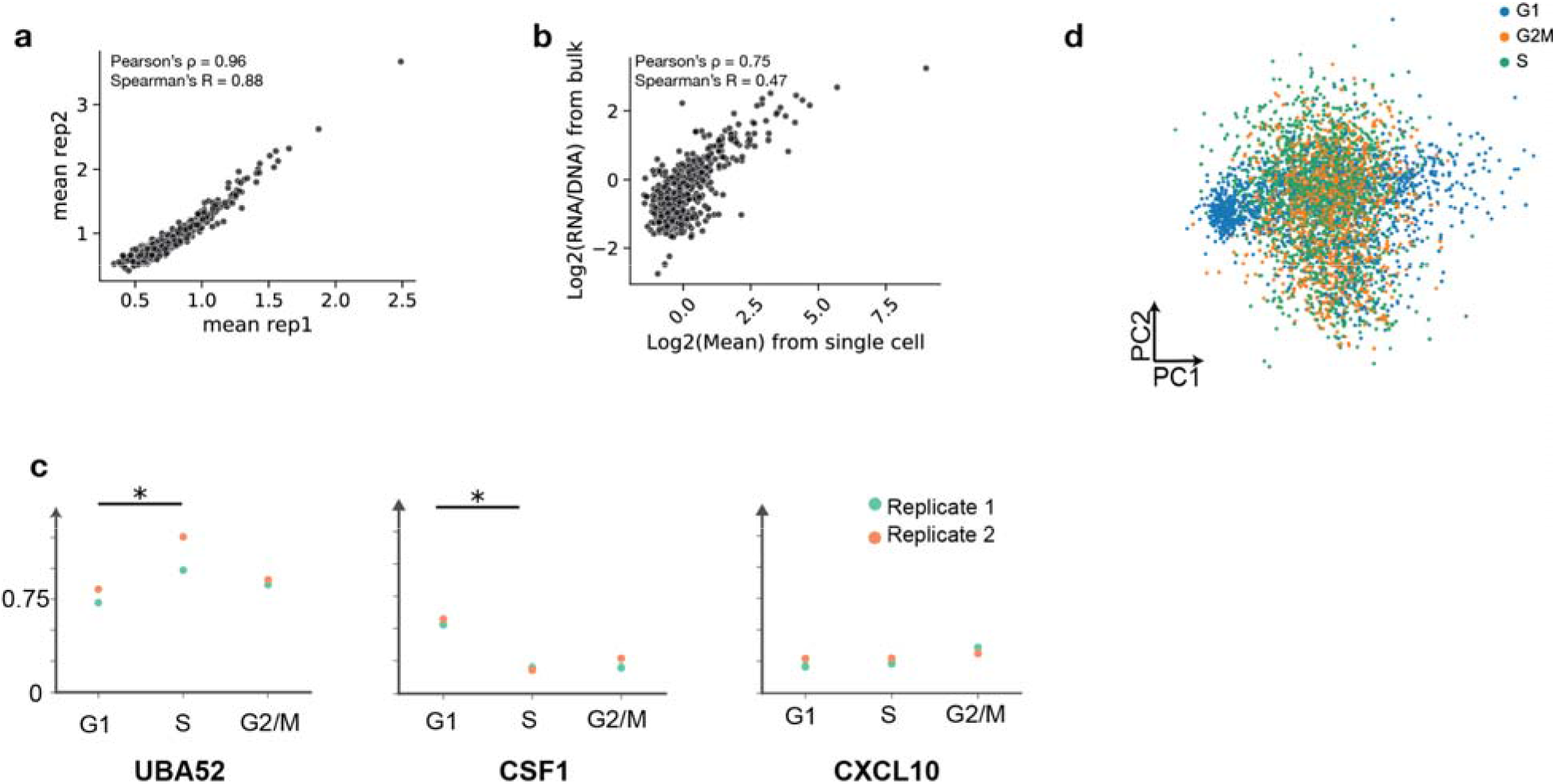
scMPRA measures CRS activity in K562 cell substates. (**a**) Reproducibility for mean expression of core promoters in K562 cells. **(b)** Correlation of bulk and scMPRA (non-UMI corrected) in K562 cells **(c)** Different dynamics of expression. For *UBA52*, the promoter is most highly expressed in S phase; whereas for *CSF1*, the promoter is most highly expressed in G1 phase. For *CXCL10*, the promoter is expressed evenly through cell cycle (Stars indicate significance from Wilcoxon rank sum test, *: p < 0.05.) **(d)** Cells no longer cluster together based on cell cycle genes after the effects of the cell cycle are removed.

**Supplementary Figure 3.**
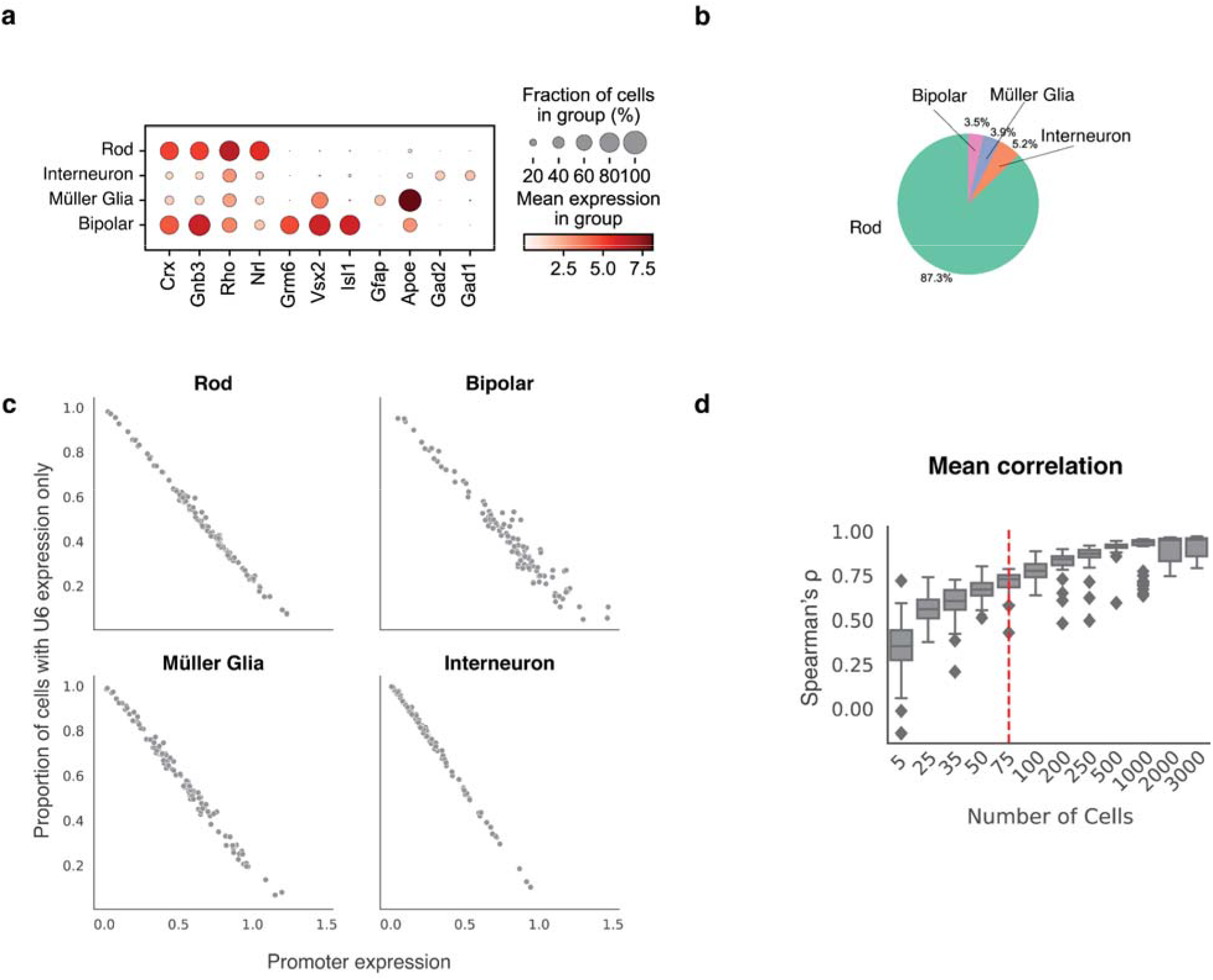
**(a)** Expression of marker genes by scRNA-seq used to identify cell types in the retina **(b)** Percentage of the total cells recovered represented by each retinal cell type **(c)** Plot showing the relationship between the mean activity of a *Gnb3* promoter variant in a given cell type (x-axis) and the proportion of cells in which that promoter variant is silent (y-axis). Individual cells in which a given *Gnb3* variant is silent are identified as cells with U6-expressed cBC, but no *Gnb3*-expressed cBC. **(d)** The correlation between biological replicates is plotted as a function of the number of cells used in the analysis.

